# *Coolpup.py:* versatile pile-up analysis of Hi-C data

**DOI:** 10.1101/586537

**Authors:** Ilya M. Flyamer, Robert S. Illingworth, Wendy A. Bickmore

## Abstract

**Motivation:** Hi-C is currently the method of choice to investigate the global 3D organisation of the genome. A major limitation of Hi-C is the sequencing depth required to robustly detect loops in the data. A popular approach used to mitigate this issue, even in single-cell Hi-C data, is genome-wide averaging (piling-up) of peaks, or other features, annotated in high-resolution datasets, to measure their prominence in less deeply sequenced data. However current tools do not provide a computationally efficient and versatile implementation of this approach.

**Results:** Here we describe *coolpup.py* – a versatile tool to perform pile-up analysis on Hi-C data. We demonstrate its utility by replicating previously published findings regarding the role of cohesin and CTCF in 3D genome organization, as well as discovering novel details of Polycomb-driven interactions. We also present a novel variation of the pile-up approach that can aid the in statistical analysis of looping interactions. We anticipate that *coolpup.py* will aid in Hi-C data analysis by allowing easy to use, versatile and efficient generation of pileups.

**Availability and implementation:** *Coolpup.py* is cross-platform, open-source and free (MIT licensed) software. Source code is available from https://github.com/Phlya/coolpuppy and it can be installed from the Python Packaging Index.

## Introduction

Major advances in the study of 3D genome organization have come from the development of a family of Chromosome Conformation Capture (3C) methods (Dekker *et al.*, 2002). While these all rely on the same principle of *in situ* proximity ligation of crosslinked and digested chromatin, the scope of each method varies depending on experimental processing and the method of quantification of the 3C library (Barutcu *et al.*, 2016). Hi-C, a genome-wide 3C-derivative, is the method of choice to investigate the organization of the whole genome (Lieberman-Aiden *et al.*, 2009; Rao *et al.*, 2014).

One of the main challenges in Hi-C remains the required sequencing depth due to the extreme complexity of good quality Hi-C libraries. The output of Hi-C is a square matrix of interactions and therefore requires a vastly greater sequencing depth than most sequencing-based approaches that simply look for enrichment of reads linearly along the genome (Lajoie *et al.*, 2014). This limits the resolution at which genomes can be analysed in 3D, since going beyond ∼5 kbp resolution requires billions of read pairs for a mammalian genome.

Looping interactions are among the most interesting features that can be studied using Hi-C. Chromatin loops bring distal regions in the genome into close proximity and are manifest in Hi-C data as foci of increased interaction frequency (Rao *et al.*, 2014). The majority of loops identified in Hi-C data from mammalian cells correspond to CTCF/cohesin associated interactions, created by loop extrusion (Fudenberg *et al.*, 2016; Sanborn *et al.*, 2015; Gassler *et al.*, 2017). CTCF/cohesin associated loops are closely related to topologically-associating domains (TADs), which in most cases are encompassed in a loop, and which can in turn contain loops. TADs have been reported to constrain enhancer-promoter communication in some cases (Lupiáñez *et al.*, 2015; Franke *et al.*, 2016; Williamson *et al.*, 2019) and might be related to genome stability (Canela *et al.*, 2017, 2019, 201), while some loops have been suggested to correspond to enhancer-promoter contacts (Rao *et al.*, 2014). In addition, distal polycomb sites can be brought together in ‘loops’ (Joshi *et al.*, 2015; McLaughlin *et al.*, 2019).

To our knowledge, currently the only robust method to identify loops *de novo* requires very deep Hi-C libraries, on the order of over a billion Hi-C contacts (Rao *et al.*, 2014). This means that the vast majority of Hi-C datasets cannot be used to identify loops. However, they can be used to quantify the average loop strength (i.e. enrichment of contacts in those loops relative to their local background). To do this one can average (or “pile up”) all areas of the Hi-C maps containing loops, that have been annotated in a high-depth dataset (Rao *et al.*, 2014). This idea is very similar to “metaplots” used, for example, in chromatin immunoprecipitation and sequencing (ChIP-seq) analysis to quantify signal in a subset of regions, except in Hi-C this is for a 2D matrix instead of a linear track. The same approach can, of course, be applied directly to the data where the loops were annotated for quantification of their average prominence. Apart from quantifying the strength of known features, the same analysis can be used to investigate whether certain regions, defined for example based on ChIP-seq peaks, tend to interact with each other on average above background. To our knowledge, the first application of pile-up-like analysis was to investigate clustering of pluripotency factor binding sites in mouse embryonic stem (ES) cells (de Wit *et al.*, 2013). Pile-up analysis can alsod aid in the discovery of novel drivers of interactions.

Hi-C is a cell population-based method, and only provides population average measurements. Several single-cell Hi-C approaches have been published (Nagano *et al.*, 2017, 2013; Stevens *et al.*, 2017; Flyamer *et al.*, 2017; Tan *et al.*, 2018), (reviewed in Ulianov *et al.*, 2017), however none of these provides data depth or resolution comparable to that which can be obtained from a population of thousands of cells (Díaz *et al.*, 2018): the resulting matrices are too sparse to analyse individual regions and only aggregate genome-wide metrics can be efficiently employed. Approaches to analyse strength of loops, TADs and genome compartmentalization from such data genome-wide have been developed (Flyamer *et al.*, 2017). These are all based on the “pile-up” approach described above using data from single cells for the regions corresponding to specific features identified in population Hi-C data, to boost the amount of reads used in the analysis.

Since its inception in the current form (Rao *et al.*, 2014), originally termed APA (“Aggregate Peak Analysis”), pile-up analysis has been used in numerous publications, both to analyse single-cell Hi-C data (Flyamer *et al.*, 2017; Gassler *et al.*, 2017; Nagano *et al.*, 2017) and as a general way of quantifying feature strength (Fudenberg *et al.*, 2016; Schwarzer *et al.*, 2017; Bonev *et al.*, 2017; Nora *et al.*, 2017; McLaughlin *et al.*, 2019; Rao *et al.*, 2017; Díaz *et al.*, 2018; Kruse *et al.*, 2019; Abdennur *et al.*, 2018; Rowley *et al.*, 2019; Krietenstein *et al.*, 2019; Hsieh *et al.*, 2019). A visual interactive tool to semi-manually classify and pile-up predefined regions has also been developed (Lekschas *et al.*, 2018). However no single computational tool can perform all the various kinds of pile-up analyses that have been used in the literature, including local and rescaled (features of different size or shape are averaged, e.g. average TADs) and off-diagonal (e.g. average loops) pile-ups with different normalization strategies (Table 1). At the same time, performing detailed analysis of Hi-C data remains difficult for non-specialists due to the absence of easy to use tools.

**Table 1:** Comparison of tools to perform pileup analysis on Hi-C data.

| Feature | Juicer | HiCExplorer | GENOVA | coolpup.py |
| --- | --- | --- | --- | --- |
| Aggregate loops | + | - | + | + |
| Aggregate region pairs | - | + | + | + |
| Interactions between two region sets | - | + | - | + |
| Local pileups | - | + | - | + |
| (Local) rescaled pileups | - | - | + | + |
| Distance normalization | - | Expected (and z-score) | Fixed shifts (for pairwise analysis) | Expected or random shifts |
| Coverage normalization | - | - | - | + |
| Anchored pileups/loop-ability | - | - | - | + |
| CLI | + | + | - | + |
| Simple text output of pileups | + | + | - | + |
| Hi-C file format | .hic | .cool, .h5, other? | HiC-pro | .cool |

Comparison of four tools for pileup analysis across a set of features: Juicer Aggregate Peak Analysis (APA) (Rao *et al.*, 2014), HiCExplorer (hicAggregateContacts and hicAverageRegions) (Ramírez *et al.*, 2018), GENOVA (APA, ATA and PE-SCAN) (Weide, 2019) and coolpup.py.

Here we present a unified command-line interface tool written in **Py**thon to **p**ile-**up** Hi-C data stored in the widely used and versatile .**cool** format (Abdennur and Mirny, 2019) (*coolpup.py*). A simple script for plotting the output of *coolpup.py* is provided in the package (*plotpup.py*), although for higher flexibility we suggest directly using *matplotlib* or another library.

Here we have applied *coolpup.py* to published data to investigate the effect of different normalization strategies on the resulting pileups, and to replicate published results to verify *coolpup.py*’s algorithm. We also present a novel variation of the pileup approach implemented in *coolpup.py* that retains some of the locus-specific information and would allow more detailed statistical analysis of looping interactions in Hi-C data. Using published single-cell Hi-C, we also investigate the dynamics of polycomb-associated looping revealing a different dynamics of looping across the cell cycle compared with CTCF loops.

## Materials and methods

### Sources of datasets & data analysis

As a proof of principal, we applied *coolpup.py* to publicly available Hi-C data (Bonev *et al.*, 2017; Nora *et al.*, 2017) using *distiller* (https://github.com/mirnylab/distiller-nf) to obtain .*cool* files filtered with a map quality (mapq) of ≥30. We used this data at 5 kbp resolution. In addition .*cool* files for single-nucleus Hi-C (snHi-C), together with coordinates of loops and TADs used in the original publication (kindly shared by Hugo Brandão) (Gassler *et al.*, 2017; Rao *et al.*, 2014), were reanalysed at 10 kbp resolution (without balancing and with coverage normalization and 10 random shifts). We also used single-cell Hi-C data for diploid mouse ES cells grown in serum from (Nagano *et al.*, 2017) (.*cool* files were kindly shared by Aleksandra Galitsyna) at 5 kb resolution. We created pile-ups for each cell in the same manner as for snHi-C, and used the average value of interactions in the central 3×3 pixel square to get the level of interaction enrichment. RING1B and H3K27me3 ChIP-seq peaks (ref. Illingworth *et al.*, 2015) were lifted over to the mm9 mouse genome assembly. The coordinates of biochemically defined CpG islands were taken from (ref. Illingworth *et al.*, 2010). CTCF ChIP-seq peaks were taken from (ref. Bonev *et al.*, 2017) and, following liftOver to the *mm9* assembly, intersected with CTCF motifs found in the mm9 genome using Biopython’s *motifs* module (Cock *et al.*, 2009). A human CTCF position-frequency matrix was downloaded from JASPAR (MA0139.1). We used only motifs with a score >7, and discounted peaks containing >1 motif.

Regions of high insulation (meaning low number of contacts crossing this regions) in the Bonev et al. Hi-C data were called using *cooltools diamond-insulation* from 25 kbp resolution data and a window size of 1 Mb. The output was filtered to exclude boundaries with strength <0.1, and then pairs of consecutive boundaries were combined to create an annotation of TADs. TADs longer than 1,500 kbp were not used due to their likely artefactual nature (based on both visual inspection, and the fact that TAD sizes are reported to be on the order of a few hundred kbp in mammalian cells (Rao *et al.*, 2014)). The same loop annotations for mouse embryonic stem (ES) cells were used as in our recent preprint (McLaughlin *et al.*, 2019).

All figure panels were created using *matplotlib (Hunter, 2007)* and assembled in Inkscape.

### *Coolpup.py* implementation

*Coolpup.py* is a versatile tool that uses .*cool* files as the main input together with a bed (chrom, start, end) or pairbed (chrom1, start1, end1, chrom2, start2, end2) file to define the regions under investigation. The tool is implemented as a *python* package which parses all arguments via *argparse*, performs the computation and saves the output file(s). It leverages the scientific python environment, taking advantage of *numpy* (Walt *et al.*, 2011), *scipy* (Jones *et al.*, 2001) and *pandas (McKinney, 2010)*. A separate CLI tool included in the package (*plotpup.py*) can be used to visualize the results, and uses *matplotlib* (Hunter, 2007). The code is available on github (https://github.com/Phlya/coolpuppy) and the package can be installed using *pip*, which then makes *coolpup.py* and *plotpup.py* available in the command line. Alternatively, all main functions can be accessed directly from *python*.

The overall procedure for piling up a lot of small regions is the following. To minimize the number of file reads (at the cost of required computer memory), a sparse representation of each chromosome’s Hi-C contact matrix is loaded into memory. Then, using an iterator, each required location (on- or off-diagonal) is individually retrieved to generate a corresponding submatrix from the data (with some specified padding around the centre of the ROI), and added to the matrix of the same shape, initialized with zeros, while keeping track of the number of summed up regions. If specified, coverage of the window on each side is recorded. Similarly, if needed, the window (and the coverage) is rescaled to a required shape. This is done for all chromosomes (optionally, in parallel using *multiprocessing*), and then all of the results are summed and then divided by the total number of windows. If specified, coverage normalization is applied at this stage. Then, unless otherwise specified, a normalization to remove the distance-dependency of contact probability is applied. In most cases the best and most efficient way is to use a (chromosome-wide) expected value for each diagonal of the matrix, which can be obtained for a cooler file using, for example, *cooltools compute-expected*. This information is then used to create a matrix of expected interactions, under the assumption that the probability of interactions only depends on distance. Alternatively, if the expected values are not available, for example for single-cell Hi-C data, this normalization can be performed using randomly shifted control regions. In that case the whole pile-up procedure is repeated, but the coordinates are randomly shifted. In the end, the resulting matrix of averaged interactions is divided by the normalizing matrix to remove effects of distance.

If not specified, balanced data with chromosome-wide expected normalization were used when creating pileups, except for the zygote and single-cell Hi-C datasets, where randomly shifted controls and coverage normalization were used instead. For the single-cell Hi-C (Nagano *et al.*, 2017) analysis we only used pairs of RING1B and convergent CTCF peaks within 100-800 kbp of each other, since previous analysis (data not shown) indicated this as the distance range where both looping modes are observed. For plotting average TADs, apart from observed/expected pileups in Figure 2B, we show the matrices after re-introduction of slow decay of interaction probability with distance. We simply multiply the observed/expected matrix by a matrix of the same shape where each diagonal *i* has the value *i*^*-0.25*^, starting from 0^0.25^=1.

**Figure 1.**
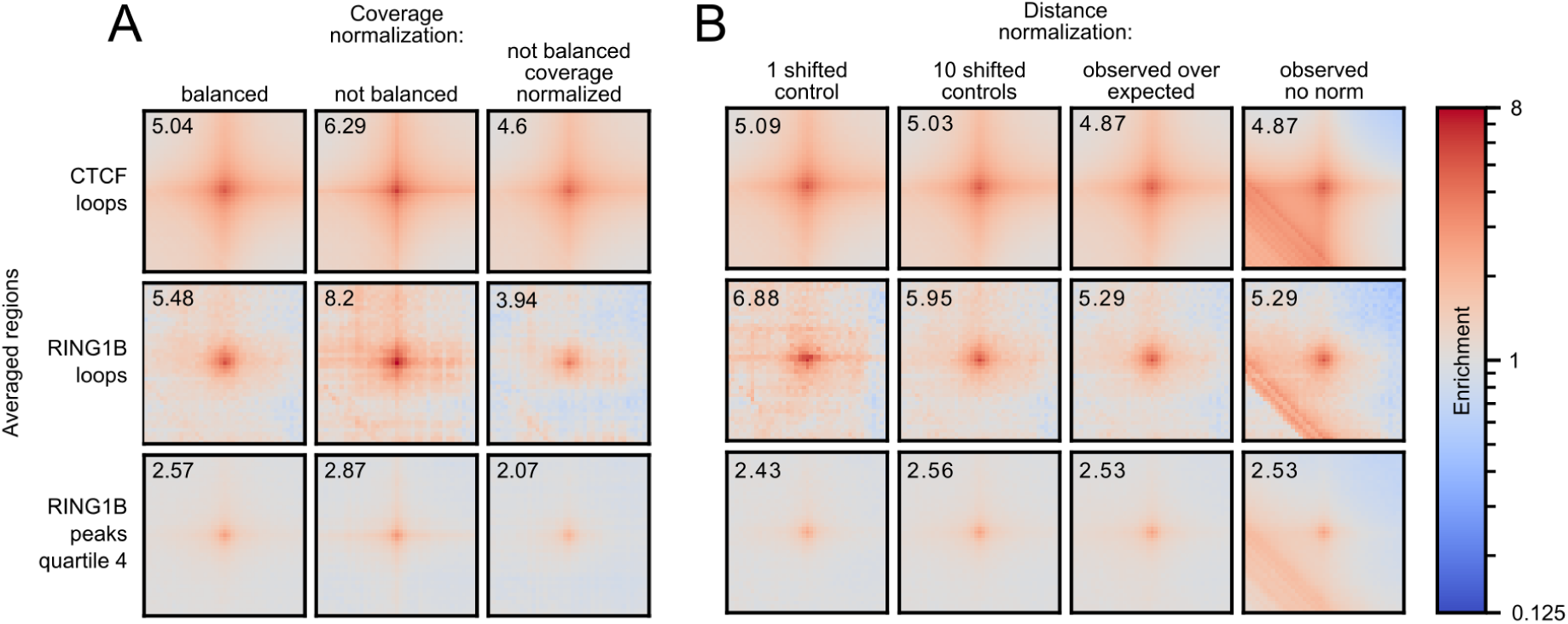
Hi-C data normalization strategies. (**A**) Comparison of coverage normalization strategies for pile-up analyses using mouse ES cell Hi-C data (Bonev *et al.*, 2017). Normalization approaches are in columns: matrix balancing (iterative correction); no normalization; no balancing with coverage normalization of the pileups. The different averaged regions are shown in rows: loops associated with CTCF (n=6536), loops associated with RING1B (n=104) (see Methods), all pairwise combinations of high RING1B peak regions from the 4^th^ quartile (by RING1B ChIP-seq read count) (n=2660 of peak regions). All pileups produced with 10 randomly shifted controls to remove short distance artefacts. All pileups are normalized to the average of the top-left and bottom-right corner pixels to bring them to same scale. Number in top left corner is value of the central pixel. 5 kb resolution with 100 kb padding around the central pixel. Colour is shown in log-scale and shows enrichment of interactions. (**B**) Same as A, but for different approaches to remove distance-dependency of contact probability with balanced data. In columns: single randomly shifted control regions per ROI; ten randomly shifted control per ROI; normalization to chromosome-wide expected; no normalization. Same rows as in (A).

**Figure 2.**
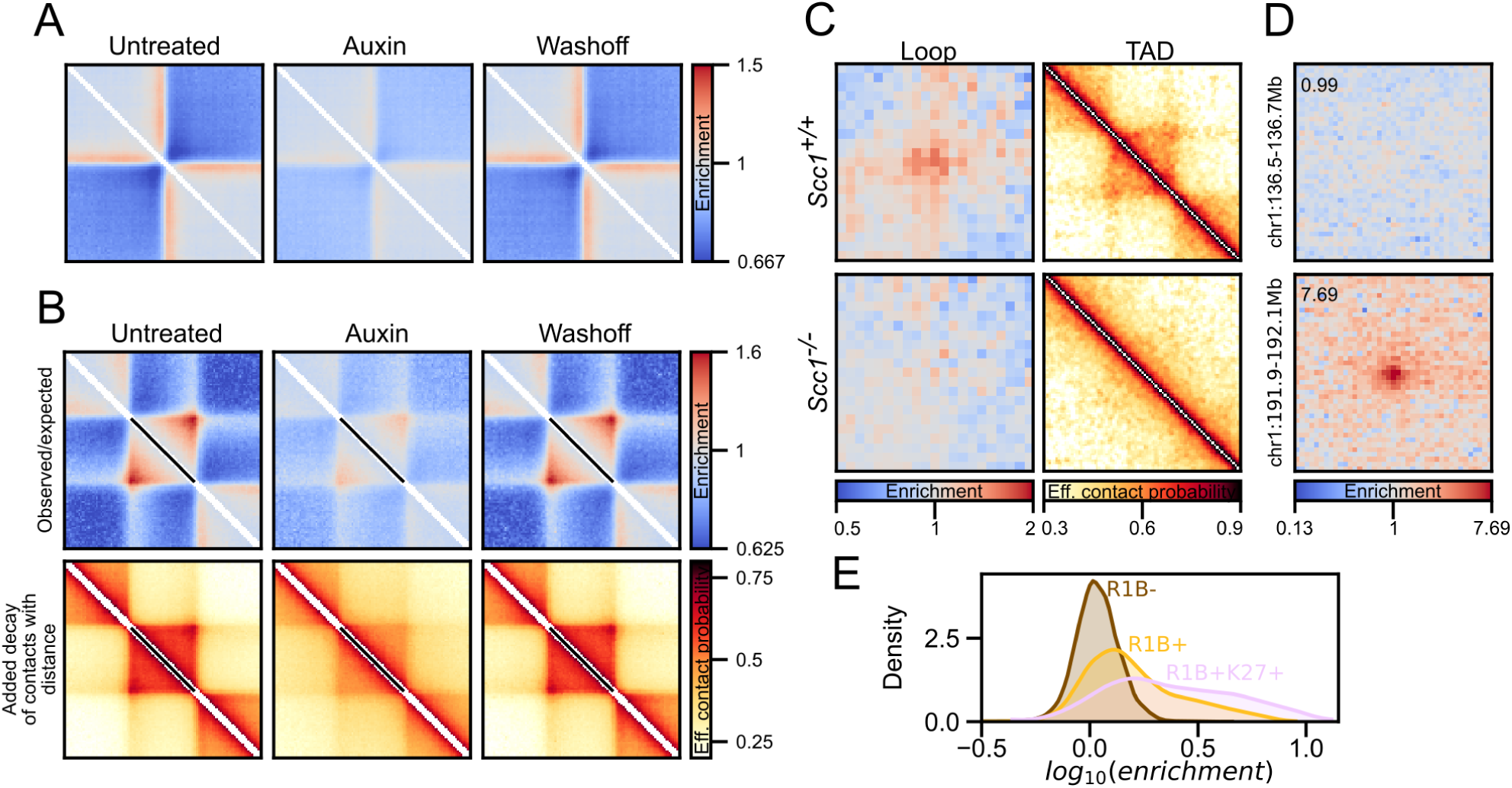
Pileup variations. (**A**) Local pileups of high insulating regions in ES cells across untreated, auxin-treated and wash-off conditions in CTCF-AID Hi-C data (Nora *et al.*, 2017). 25 kb resolution data with 1,000 kb padding around the central pixel. (**B**) Local rescaled pileups of TADs (defined based on high insulating regions) across same data as in (A) from 5 kbp resolution data. Top row: raw observed over expected pileups; bottom row: pileups after artificial re-introduction of shallow decay of contact probability (P_c_=s^-0.25^, see Methods) for easier visual interpretation of the data (Flyamer *et al.*, 2017). (**C**) Loop and rescaled TAD pileups for pooled single-cell Hi-C data showing loss of structures in Scc1^-/-^ zygotes (Gassler *et al.*, 2017) (**D**) Two examples of anchored pileups from RING1B+/H3K27me3+ CpG islands, with no visible enrichment (top), or with very prominent enrichment (bottom). The anchored region is on the left side of the pileup, and its coordinates (including the padding) are shown on the left. The value of the central pixel (“loopability”) shown in top left corner. (**E**) Distribution of “loopability” values of CpG islands not bound by RING1B, CpG islands bound by RING1B, and CpG islands bound by RING1B and also marked by H3K27me3.

### Performance profiling

*Coolpup.py* performance was tested on the University of Edinburgh Open Grid Scheduler cluster (Eddie3). We used the Hi-C datasets for mouse ES cells (Bonev *et al.*, 2017; Nora *et al.*, 2017) for testing. To generate the required large number of coordinates for testing, we used coordinates of the B3 repeat from RepeatMasker track available from UCSC Genome Browser. For coordinate pairs, we used all pairs of convergent CTCF sites, described above. A separate job was submitted for each measurement, and the runtime of the *coolpup.py* call was recorded. Subsets of different sizes were generated using *coolpup.py*’s `--subset` argument. Where not specified, 1 compute core was utilized. The same procedure was performed for HiCExplorer *hicAggregateContacts*, except *shuf* was used to generate a random sample of required size. The following arguments were also provided to mimic *coolpup.py* behaviour as close as possible: *--range 105000:1000000000000 --avgType mean --transform obs/exp*. All measurements were performed 5 times. Plotted in Figure 4 are actual measured runtime values, the line shows mean values and shaded area - ±95% confidence interval, using the *seaborn* plotting package (Michael Waskom *et al.*, 2018).

**Figure 3.**
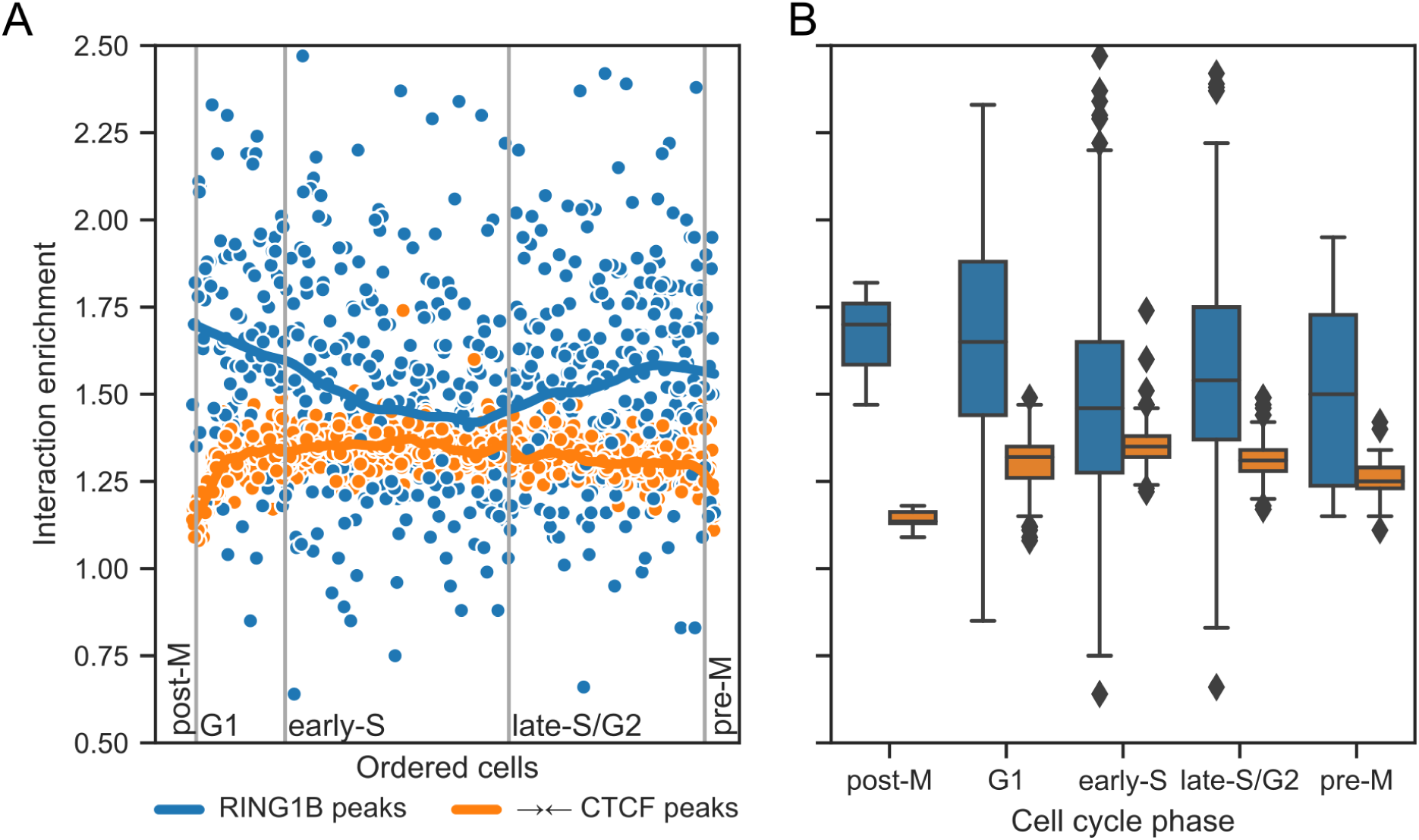
Chromatin looping dynamics across cell cycle. **(A)** Hi-C interaction enrichment levels for single cells ordered along the cell cycle (Nagano *et al.*, 2017) for CTCF- and RING1B-associated loops. Curves represent LOWESS-smoothed data for easier interpretation. **(B)** Distribution of enrichment values in all cell cycle stages from data in (A).

**Figure 4.**
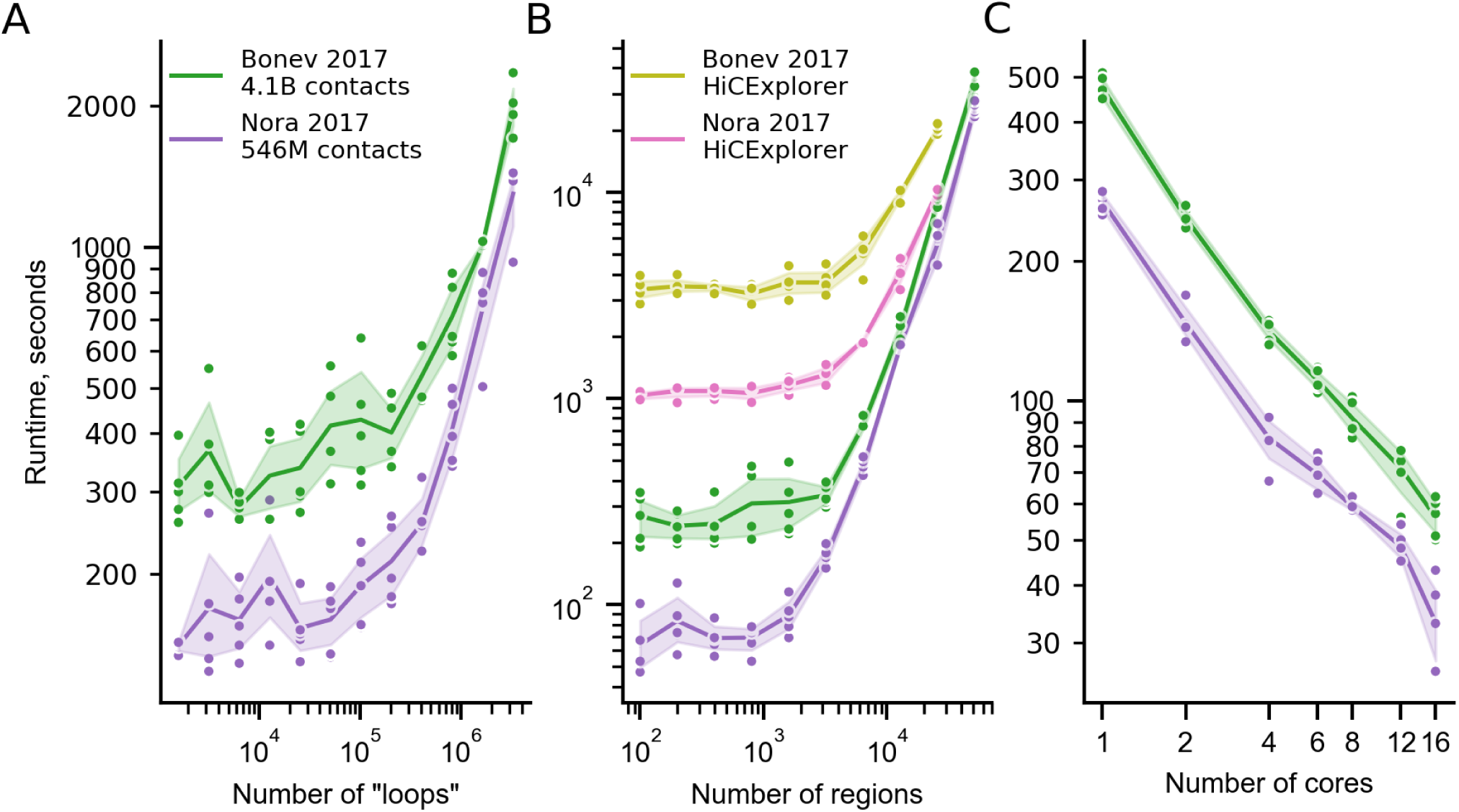
Performance profiling. (**A**) Runtime (seconds) of *coolpup.py* with varying number of averaged “loops” for two Hi-C datasets with different depth. (**B**) Same as (A), but for number of linear regions between which interactions are averaged. Also shown is runtime for HiCExplorer *hicAggregateContacts*. Note that the longest timepoint for HiCExplorer required over 512 Gb RAM and was not computed. (**C**) Runtime of the same analysis with 5000 linear regions and a varying number of cores.

## Results

### Different normalization strategies implemented in *coolpup.py*

Hi-C data can be normalized in different ways to remove either technical biases, or uninteresting (in this context) biological signal of the decay of contact probability with genomic distance. *Coolpup.py* provides ways to deal with both of these problems.

Hi-C data are usually normalized to remove systematic biases, such as GC-content or restriction site frequency (Yaffe and Tanay, 2011). *Cooler* implements a matrix balancing (visibility equalization) approach to remove all potential biases (Imakaev *et al.*, 2012) and, when available, it is recommended to use balanced data for pile-ups. However sometimes, for example in single-cell Hi-C, removing biases is impossible due to the sparsity of data. Therefore, using unbalanced data is also an option in *coolpup.py*. However, because of the averaging of multiple regions during the pile-up procedure, the effect of biases can be partially mitigated by normalizing the matrix by the coverage (i.e. the total number of contacts of the bins in the chromosome) of the averaged regions (Flyamer *et al.*, 2017). As illustrated in Figure 1A for both CTCF and polycomb (RING1B) loops, this approach reduces coverage variability between bins and removes sharp crosses from the central bin that is present with unbalanced data. This normalization seems to slightly over-correct, i.e. the value of the central pixel is consistently somewhat lower than when using balanced data. However, the results overall look more similar to balanced data than without coverage normalization.

In addition to normalization to remove biases, it is often desirable to remove the distance-dependency of contact probability in Hi-C data, since it can have a very strong effect on the resulting pileup by introducing artefacts from very short distances, and sometimes can obscure interesting properties, such as enrichment in the centre of the pileup. However it is worth bearing in mind that this normalization can also hide real signal in the data, such as enrichment of interactions in the lower left corner, observed for CTCF-anchored loops (data not shown). The general approach to perform this normalization is to create a vector of expected contact frequency, which usually corresponds to the averaged value of the Hi-C map at each diagonal per chromosome. A file with such information (for balanced data) can be obtained using *cooltools compute-expected* then used in *coolpup.py* to normalize the pileups. However sometimes the expected information is unavailable, for example, in single-cell Hi-C it can be too noisy. In that case, an alternative approach to remove distance-dependency of contact frequency can be used: for each position in the Hi-C map being averaged, a matched set of randomly shifted control regions with the same distance separation is used (Flyamer *et al.*, 2017). In this way, by creating many such control regions for each region of interest (ROI), it is possible to estimate the expected frequency of interactions even for sparse single-cell Hi-C data. As shown in Figure 1B, both of these approaches are excellent at removing artefacts resulting from short-range interactions in the pileups and produce visually indistinguishable results. However, for a small set of regions (e.g. RING1B associated loops) a higher number of randomly shifted controls for each ROI is required to prevent noise. We note that for local pile-ups (especially with rescaling; see below) random controls perform better than simple normalization to expected values (data not shown).

### Applications of pile-ups

As well as the basic pile-up procedure, there are multiple variations built in to *coolpup.py* which are tailored to answer different biological questions. The following ones are trivial, but worth mentioning. For example, often it is desirable to restrict the minimal and/or maximal separation of analysed sites, either to remove short-range artefacts, or to analyse the distribution of enrichment signal across different distance scales. Only certain chromosomes might need to be included, or, with too many regions of interest, a random subset can be taken to speed up the computation.

A popular variation of the pile-up approach is “local” pile-up: an analysis which focuses on near-diagonal features. For example, we averaged regions of high insulation annotated in the deep ES cell Hi-C dataset to visualize insulation strength after Auxin-induced degradation of CTCF (Nora *et al.*, 2017) (Figure 2A). In this case the pileups are performed in the same way as previous off-diagonal pileups, however the regions that are averaged lie on the main diagonal of the Hi-C map. A variation of this approach is local pileups with rescaling to analyse features of different size, for example, TADs (Flyamer *et al.*, 2017). As an example, TADs, based on aforementioned regions of high insulation annotated in data from (ref. Bonev *et al.*, 2017), were averaged to visualize changes in local interaction strength upon CTCF degradation (Nora *et al.*, 2017) (Figure 2B). Here all windows centred on regions of interest are rescaled to the same size, and then averaged.

Pileups are a particularly important approach to analysing very low depth datasets to uncover genome-wide average patterns, which are indiscernible when looking at individual regions in such shallow data. Here we apply *coolpup.py* to reproduce results from a dataset comprising pooled data from a few single cells, to show a loss of loops and TADs in mouse zygotes lacking SCC1 (RAD21), the kleisin subunit of cohesin (Gassler *et al.*, 2017). Since the material is so limiting and data are based on single cells, the total number of contacts in this dataset is very low: 4.8 and 9.2 million contacts in *Scc1*^+/+^ and *Scc1*^-/-^, respectively. However, we successfully performed pileups, both with “traditional” averaging of loops, and local pileups of TADs with rescaling, and observe the loss of both loops and TADs upon deletion of cohesin, comparable to the original study (Figure 2C).

All pile-up approaches include averaging of multiple regions, a drawback of which is loss of locus-specific information. We therefore designed a novel approach that retains some information about the specific loci used in the analysis. In this approach, we pile-up a single region against multiple other regions; the same can be done for each of many regions in a set against all other regions. Then by extracting the value in the central pixel in pileups for each region, we can get a “loop-ability” value, which can then be related to other features of analysed regions, such as the level of occupancy by different factors. To confirm that this approach can work, we checked some example regions that displayed high or low level of “loop-ability”, to ensure that the values we observed were not due to noise from piling up interactions of a single region (see two examples in Figure 2D). A simple proof of principle analysis highlights the interactions between sites bound by polycomb group proteins in mouse ES cells (data from Bonev *et al.*, 2017). By splitting the CpG islands (data from Illingworth *et al.*, 2010) - the main targets of polycomb binding in ES cells - in the mouse genome into RING1B (a core component of Polycomb Repressive Complex 1 - PRC1) negative, RING1B positive, and RING1B and H3K27me3 positive sets (data from Illingworth *et al.*, 2015), we observe high “loop-ability” values for the two latter groups, while the RING1B negative CpG islands have close to no enrichment (Figure 2E).

Pileups are an invaluable tool when analysing Hi-C data from single cells, since averaging features across the whole genome helps to circumvent the sparsity of the data. Here we apply *coolpup.py* to analyse the looping interactions across the cell cycle using a published single-cell Hi-C dataset from hundreds of mouse ES cells (Nagano *et al.*, 2017). We compared the enrichment of interactions in different cell cycles stages for CTCF- and RING1B-associated interactions (see Figure 3A, 3B). For convergent CTCF sites, we detected the loss of loop strength in early G1, and in pre- and post-mitotic cells, consistent with the original publication (Nagano *et al.*, 2017).

In contrast, the interactions between RING1B binding sites have a very different dynamic across the cell cycle. They are at their weakest during S phase, progressively strengthening during G2 and not reaching their peak until early G1. This is consistent with the cell cycle kinetics of H3K27me3 abundance at polycomb marked sites with H3K27me3 levels lowest during S phase where they are diluted after the replication fork, and levels of H3K27me3 only accumulating slowly through G2 and not peaking again until G1 of the next cell cycle (Reverón-Gómez *et al.*, 2018).

### *Coolpup.py* can deal with huge numbers of regions

Creating pileups from intersections of genomic regions can require averaging a huge number of 2D windows: the number of 2-combinations grows quickly with the number of regions. For example, with ∼1000 regions per chromosome (which is approximately equivalent to the number of genes), requires averaging of ∼10,000,000 regions for the whole genome, several orders of magnitude more than the number of regions usually averaged, such as number of annotated loops (∼10,000). Therefore, it is important for a general-purpose tool for creating pileups to scale well with the number of averaged 2D windows. To facilitate this, *coolpup.py* performs a very low number of read operations on the Hi-C data – only once per chromosome (or twice, when using randomly shifted controls). Whilst this necessitates that the whole Hi-C matrix of a chromosome has to be loaded into memory, it is only stored in a sparse format, and so conventional Hi-C datasets can be analysed on a regular desktop (although multi-billion contact datasets might require a high-memory machine; data not shown).

To test the performance of *coolpup.py* and how this depends on number of regions of interest, we measured the runtime with varying number of two-sided coordinate pairs (mimicking loop annotation) (Figure 4A), and varying the number of one-sided coordinate interactions being averaged (Figure 4B). We used both deep (Bonev *et al.*, 2017), and “regular depth” Hi-C data (Nora *et al.*, 2017) from mouse ES cells. With both datasets, the runtime was almost constant up to a certain number of “loops” (∼1-2×10^5^), where it starts quickly increasing (Figure 4A). Notably, the best annotations that exist to date only contain <40,000 loops (Krietenstein *et al.*, 2019), and therefore this would fall within the flat part of the curve. Similarly, in the latter analysis, runtime didn’t increase up to 1600 and 3200 regions of interest for the Nora et al. and Bonev et al. datasets, respectively. Importantly, in both analyses the difference in time between datasets with almost 10-fold sequencing depth difference is not very large, and probably largely driven by differences in time required to read the data from disk. When similar analysis was performed using HiCExplorer *hicAggregateContacts*, the runtime was >10-fold longer for each dataset with low numbers of regions (Figure 4B), and the analysis required much more memory since the algorithm uses dense data structures and stores each submatrix of interest in memory (required >100 Gb for the Bonev et al. dataset; the longest time-point required over 512 Gb of RAM and was not computed, while *coolpup.py* only needed ∼8 Gb for any calculation). HiCExplorer implementation computes observed/expected matrix for every calculation and can’t use precomputed expected values, which at least partially accounts for much longer runtimes.

Since *coolpup.py* supports parallel processing to speed up analyses, we also tested how well it scales with the number of computer cores used. We measured the runtime of the same analysis performed with varying number of cores (Figure 4C) and showed that the runtime shortened linearly with additional processes. This means the parallelization strategy used in *coolpup.py* is efficiently utilizing available CPU cores and when available, we recommend using many cores to speed up computation, although this would also significantly increase memory requirements.

## Discussion

With the large efforts being made in deciphering the structure and function of the genome in 3D, efficient, robust and versatile tools are required to facilitate quick hypothesis testing. Unlike for RNA-seq, ChIP-seq and other genome-wide methods, analysis of complex Hi-C data remains a challenge only readily accessible to specialists in the field due to an absence of easy to use informatics tools, with a few exceptions. One popular analysis applied to Hi-C data is pile-ups, which show an average genome-wide view of a selected set of regions in the 2D Hi-C interaction matrix: a very visual and intuitive approach to analysing data.

Here we presented *coolpup.py*, a versatile tool to perform pile-up analysis on Hi-C data in .*cool* format. Apart from simple generation of pile-ups, *coolpup.py* can be used to explore different data normalization strategies. While we recommend using balanced data with normalization to chromosome-wide expected interaction frequency, in certain cases a different normalization strategy can be beneficial. Similarly, exploring other parameters of the algorithm (such as minimal separation between averaged loop bases, or minimal length of locally averaged features) is straightforward with *coolpup.py*. Using our tool, we reproduced published results on the role of CTCF and cohesin in generating chromatin loops and TADs. We have shown application of *coolpup.py* to both low coverage Hi-C data (merged snHi-C data), and extremely sparse single-cell Hi-C data. The latter analysis not only replicated published data on CTCF-mediated looping changes across the cell cycles, but also revealed novel cell cycle dynamics of polycomb-associated interactions with highest contact enrichment around the time of mitosis. We note that these observations are generally consistent with the dilution and slow recovery of the H3K27me3 mark after the replication fork (Alabert *et al.*, 2015; Reverón-Gómez *et al.*, 2018), as well as an antagonistic relationship between cohesin-mediated loop extrusion and looping between RING1B target sites, reported previously (Rhodes *et al.*, 2019). These observations also pose a question of whether polycomb-associated interactions persist in metaphase chromosomes - a possibility since components of CBX2-containing PRC1 remain associated with metaphase chromosomes (Zhen *et al.*, 2014). These novel insights highlight the exploratory power of pile-up analysis.

Since *coolpup.py* is designed as a command-line tool and allows reading the coordinates of regions from standard input, it is compatible with computational pipelines, and can be readily used in shared computing environments. Moreover, it remains accessible for non-specialists with minimal knowledge of the command line and no programming experience. *Coolpup.py* should aid in improving reproducibility by providing a standardised approach for pile-up analysis which is intuitive and therefore accessible to both specialists and non-specialist alike. We hope that it will facilitate research into the 3D organization of the genome by allowing easy to use, versatile and efficient generation of pileups.

## Availability of data and materials

*Coolpup.py* is available on GitHub (https://github.com/Phlya/coolpuppy, doi: 10.5281/zenodo.3237784) and can be installed from the Python Package Index (PyPI). It is distributed under the permissive MIT license. All code used to generate the results and figures for this paper is available on GitHub (https://github.com/Phlya/coolpuppy_paper).

## Acknowledgements

We are grateful to Maksim Imakaev, Nezar Abdennur, Anton Goloborodko, Hugo Brandão and Sergey Venev for discussions, help and advice about the *cooler* ecosystem, and to Aleksandra Galitsyna for sharing .*cool* files for the Nagano et al., 2017 dataset. This work has made use of the resources provided by the Edinburgh Compute and Data Facility (ECDF) (http://www.ecdf.ed.ac.uk/). We also thank Sergey Ulianov for critically reading the manuscript.

## Authors’ contributions

I.M.F. designed *coolpup.py*, performed the Hi-C analysis and wrote the manuscript. R.S.I. performed ChIP-seq analysis. W.A.B. supervised the project. All authors read, contributed to and approved the final manuscript.

